# A population-averaged structural connectomic brain atlas of 422 HCP-Aging subjects

**DOI:** 10.1101/2023.06.01.543240

**Authors:** Yiming Xiao, Jonathan C. Lau, Terry Peters, Ali R. Khan

**Affiliations:** Department of Computer Science and Software Engineering, Gina Cody School of Engineering and Computer Science, Concordia University, Montreal, QC, Canada; Centre for Functional and Metabolic Mapping, Robarts Research Institute, Western University, London, ON, Canada; Department of Clinical Neurological Sciences, Division of Neurosurgery, Schulich School of Medicine & Dentistry, Western University, London, ON, Canada; Department of Medical Biophysics, Schulich School of Medicine & Dentistry, Western University, London, ON, Canada; School of Biomedical Engineering, Western University, London, ON, Canada; The Brain and Mind Institute, Western University, London, ON, Canada

**Keywords:** Magnetic resonance imaging (MRI), Diffusion MRI, Aging, Structural connectivity, Subthalamic nucleus, Atlas

## Abstract

Population-averaged brain atlases, that are represented in a standard space with anatomical labels, are instrumental tools in neurosurgical planning and the study of neurodegenerative conditions. Traditional brain atlases are primarily derived from anatomical scans and contain limited information regarding the axonal organization of the white matter. With the advance of diffusion MRI that allows the modelling of fiber orientation distribution (FOD) in the brain tissue, there is an increasing interest for a population-averaged FOD template, especially based on a large healthy aging cohort, to offer structural connectivity information for connectomic surgery and analysis of neurodegeneration. The dataset described in this article contains a set of multi-contrast structural connectomic MRI atlases, including T1w, T2w, and FOD templates, along with the associated whole brain tractograms. The templates were made using multi-contrast group-wise registration based on 3T MRIs of 422 Human Connectome Project in Aging (HCP-A) subjects. To enhance the usability, probabilistic tissue maps and segmentation of 22 subcortical structures are provided. Finally, the subthalamic nucleus shown in the atlas is parcellated into sensorimotor, limbic, and associative sub-regions based on their structural connectivity to facilitate the analysis and planning of deep brain stimulation procedures. The dataset is available on the OSF Repository: https://osf.io/p7syt.

**Specifications table:** 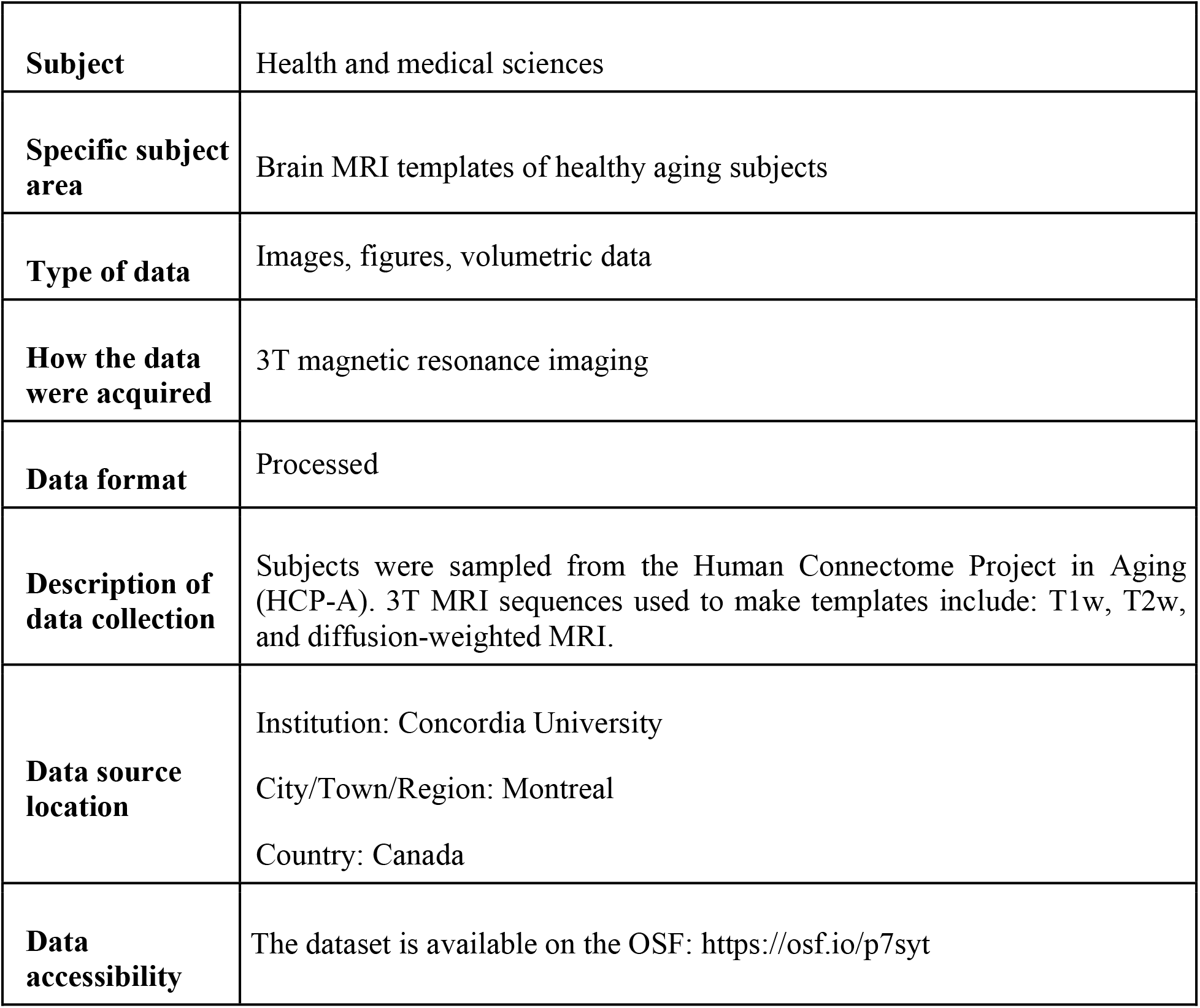

**Value of the data:**

- These publicly available templates were created using 422 HCP-Aging subjects, representing averaged anatomical and structural connectivity features of the healthy aging brain.
- Matching T1w, T2w, and fiber orientation distribution (FOD) templates are provided, together with the associated tractograms that contain 20K and 2 million streamlines.
- Segmentation of 22 subcortical structures and probabilistic tissue maps are provided to enhance the usability of the templates.
- Parcellation of the subthalamic nucleus, based on structural connectivity of the included cohort at 0.3 × 0.3 × 0.3 mm^3^ resolution, is provided.

## Data description

In neurosurgical planning and research of neurodegenerative conditions, brain atlases are crucial tools that provide the common coordinates for spatial normalization and anatomical navigation, as well as complementary physiological information (Fonov et al., 2011; Hawrylycz et al., 2012; Xiao et al., 2015; Xiao et al., 2017). While structural scans, involving T1-weighted (T1w) and T2-weighted (T2w) MRI contrasts have been traditionally used to construct brain templates, they offer limited information regarding the axonal organization of the white matter, which is of increasing interests in connectomic neurosurgery (Horn et al., 2017) and analysis of neurological conditions (Abhinav et al., 2014; Xiao et al., 2021). With recent progress in diffusion MRI (dMRI) that allows the modelling of fiber orientation distribution (FOD) in the brain tissue, there is a rising demand to incorporate the relevant information in new generations of brain atlases. Furthermore, although a few structural connectomic atlases in the forms of FOD templates or tractograms (Fan et al., 2016; Yeh et al., 2018; Aleman-Gomez et al., 2022; Lv et al., 2023) have been recently introduced, there is still a lack of population-averaged brain atlases of the same type that are derived from a large aging population with corresponding anatomical templates from the same cohort. The dataset described in this article is a collection of multi-contrast structural connectomic brain atlases, created based on 3T scans of 422 healthy subjects from the Human Connectome Project in Aging (HCP-A) group (Harms et al., 2018; Bookheimer et al., 2019). The set of atlases contain three co-registered population-averaged MRI templates. The T1w and T2w MRI templates are provided in two different resolutions: 0.5×0.5×0.5 mm^3^ and 0.3×0.3×0.3 mm^3^ while the FOD template is at 1×1×1 mm^3^ resolution. From the FOD template, two whole brain tractograms with 20K and 2 million streamlines were generated. To aid anatomical navigation and segmentation, probabilistic tissue maps and labels of 22 subcortical structures are included. For the subthalamic nucleus (STN), sensorimotor, limbic, and associative sub-regions were parcellated based on their structural connectivity and could be used to facilitate the planning of deep brain stimulation (DBS) procedures. Different components of the resulting atlases are shown in Fig. 1 to Fig. 4.

**Figure 1:**
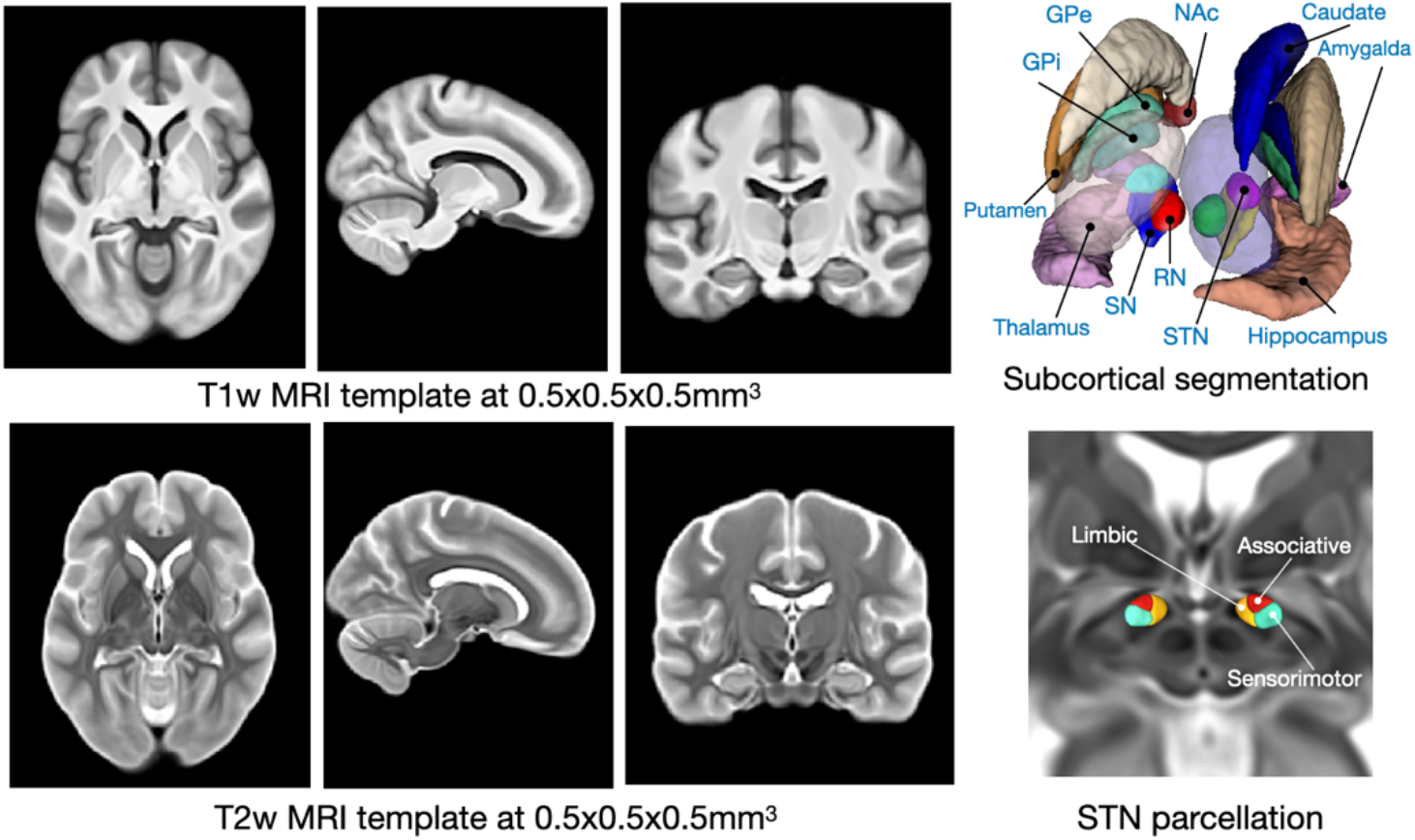
Demonstration of the T1w and T2w population-averaged MRI templates at 0.5×0.5×0.5 mm^3^ resolution, with the associated subcortical structure segmentation and structural connectivity-based parcellation of the subthalamic nucleus (STN).

**Figure 2:**
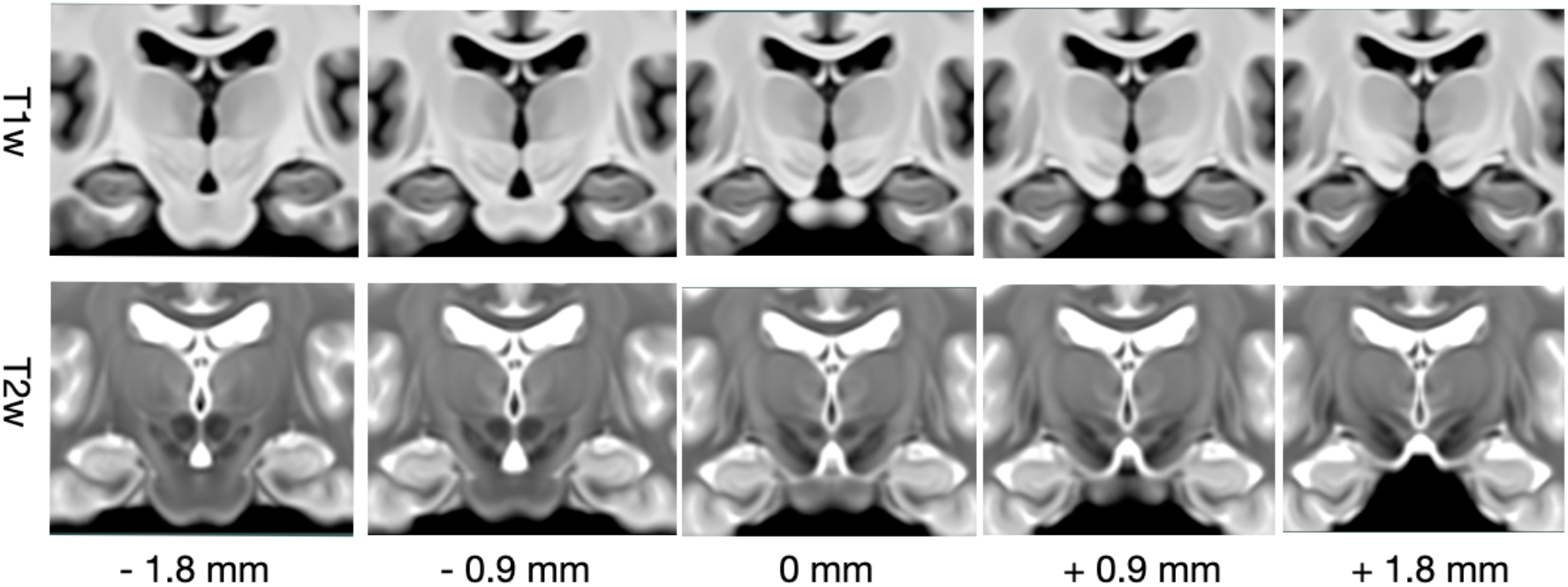
Demonstration of matching coronal views of the T1w and T2w population-averaged MRI templates at 0.3×0.3×0.3 mm^3^ resolution in the first and second row, respectively. Between two adjacent columns, a distance of 0.9mm in the coronal direction is shown, and the left to right columns show different coronal slices from the anterior to the posterior direction.

**Figure 3:**
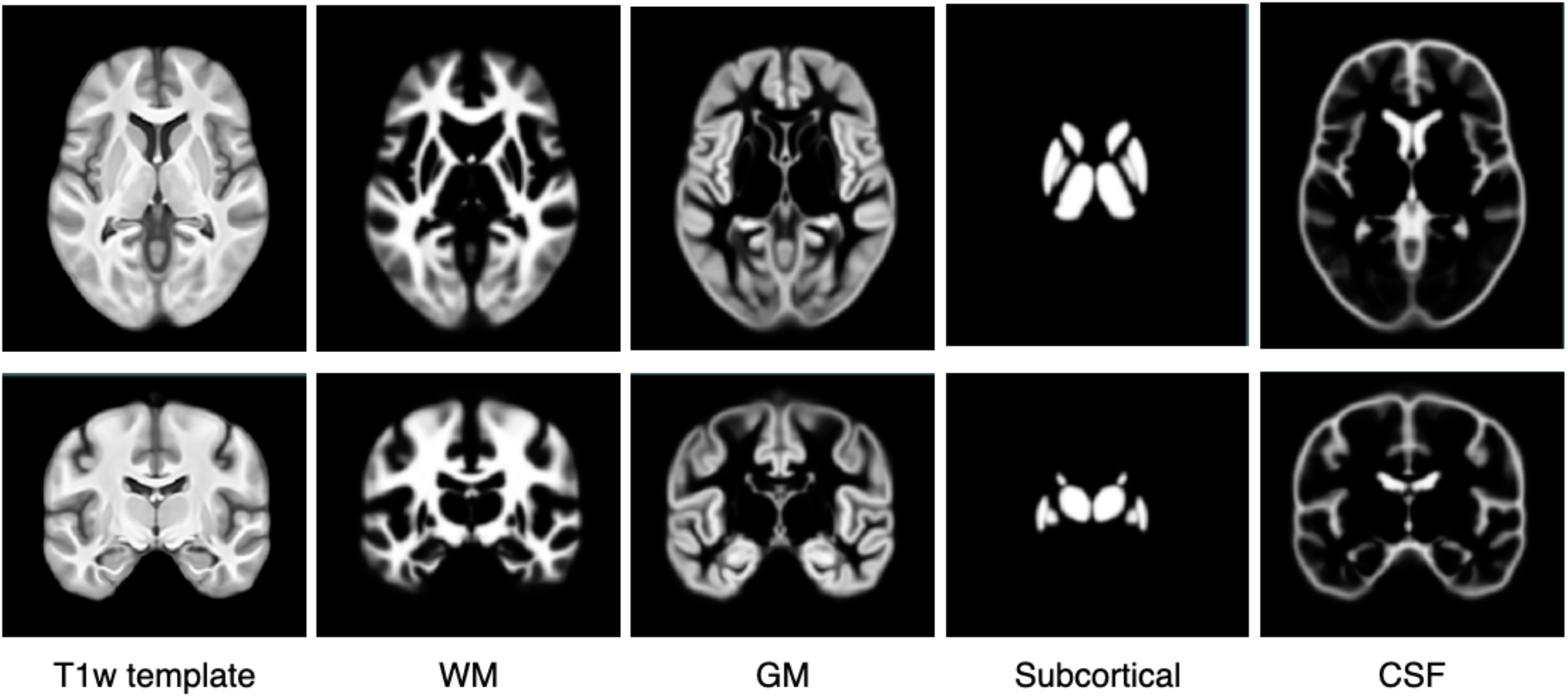
Demonstration of population-averaged probabilistic tissue segmentation maps (0.5×0.5×0.5 mm^3^ resolution) with the first and second row showing the matching axial and coronal views, respectively. From left to right: T1w template, white matter map, cortical grey matter, subcortical grey matter map, and CSF map.

**Figure 4:**
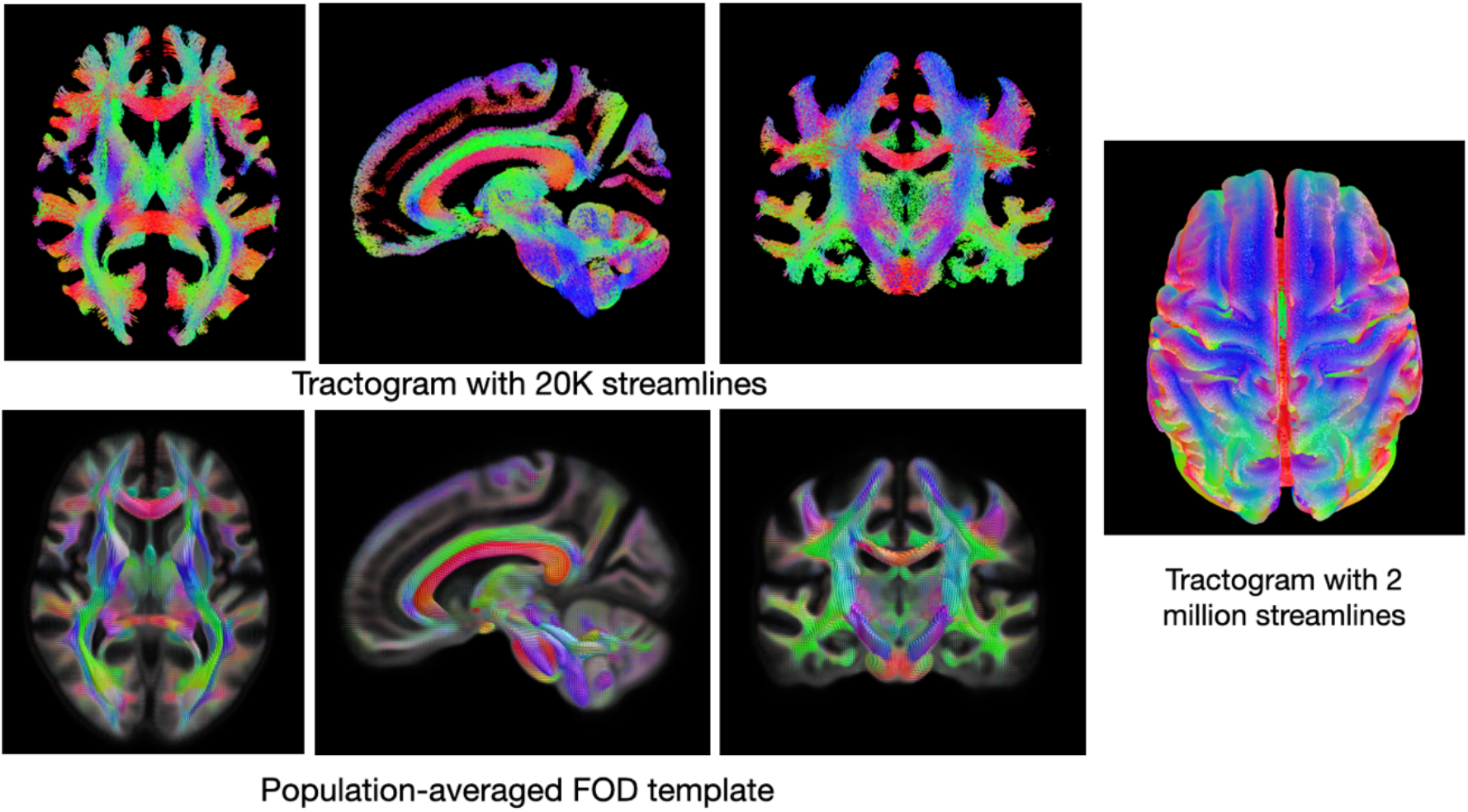
Demonstration of the population-averaged FOD template shown in the matching views of the associated tractogram with 20K streamlines, and a 3D rendering of the resulting tractogram with 2 million streamlines.

## Experimental design, materials and methods

### Subjects and image acquisition

From the Human Connectome Project in Aging (HCP-A) cohort, 436 healthy aging subjects (236 females, age=58.0±14.5 yo) were randomly sampled initially, and their T1w, T2w, and diffusion MRI scans were acquired. After image pre-processing, visual quality control was conducted by the author YX to remove subjects with poor processing outcomes (e.g., failed registration and segmentation), and 422 subjects (228 females, age= 57.5±14.4 yo) were employed to produce the final brain atlases. For each included subject, 3T T1w, T2w, and dMRI scans were obtained on Siemens Prisma 3T MRI systems with 32-channel head coils. Specifically, the T1w MRIs were acquired with a multi-echo T1w MPRAGE sequence (resolution = 0.8×0.8×0.8 mm^3^, TR/TI = 2500/1000 ms, TE = 1.8/3.6/5.4/7.2 ms, flip angle of 8 deg), and the final image was produced by the root-mean-square of the scans from all echoes. For the T2w MRI, a T2w SPACE protocol (resolution = 0.8×0.8×0.8 mm^3^, TR/TE=3200/564 ms) was used. Finally, the dMRI scans were acquired in opposite anterior-posterior phase-encoding directions with a pulsed gradient spin-echo sequence: resolution = 1.5×1.5×1.5 mm^3^, TR/TE = 3230/89.50 ms; b-values = 1500 and 3000 s/mm^2^ (92∼93 directions per shell) with 28 b-value = 0 s/mm^2^ images. More detailed image acquisition protocols of HCP-A and their differences from the HCP young adult group can be found in (Harms, et al., 2018).

### Image pre-processing

For each included subject, the T1w and T2w scans were pre-processed with non-local means denoising (Coupe et al., 2008), non-uniformity correction (Tustison et al., 2010), and intensity normalization. Then, the T2w MRI was rigidly registered to the corresponding T1w scan. The brain masks were generated using BEaST (Eskildsen et al., 2012) based on the T1w MRIs and were used to extract the brain volumes for the co-registered T1w and T2w scans. With the FAST algorithm (Zhang et al., 2001), the pre-processed T1w MRIs were segmented into probabilistic maps of white matter (WM), cortical grey matter (GM), subcortical grey matter, and cerebrospinal fluid (CSF) in the value range of [0,1].

Each dMRI scan was first processed with denoising (Veraart et al., 2016), distortion correction, unringing, eddy current correction (Andersson and Sotiropoulos, 2016), and bias field correction (Tustison, et al., 2010). The computation of individual FOD maps was conducted using the MRtrix3 software suite (Tournier et al., 2019). First, tissue-specific response functions were estimated for all subjects, from which an averaged group response function was computed for each tissue type. Then, individual FOD maps were estimated with a multi-shell, multi-tissue constrained spherical deconvolution (MSMT-CSD) algorithms (Jeurissen et al., 2014), by using the group-averaged response function (maximum spherical harmonic degree = 8). Finally, the resulting FOD maps were resampled from 1.5×1.5×1.5 mm^3^ to 1×1×1 mm^3^ for MRI template construction and tractography.

### MRI templates and tractogram construction

The unbiased population-averaged FOD template was created using multi-contrast group-wise registration jointly based on the FOD map and probabilistic tissue segmentation of the GM and CSF, using the ‘*population_template*’ script in MRtrix3. Here, fuzzy segmentations of the red nucleus (RN), substantia nigra (SN), and subthalamic nucleus (STN) were obtained for each subject by warping the labels in the MNI-PD25 atlas (Xiao, et al., 2015; Xiao, et al., 2017) with trilinear interpolation through the deformation field from atlas-to-subject nonlinear registration using the ANTs package (Tustison et al., 2014). The final probabilistic GM segmentation of each subject was generated by combining the cortical and subcortical GM segmentation maps from the FAST algorithm, as well as the fuzzy segmentation of the RN, SN, and STN. To prepare for the group-wise registration, all FOD maps were resampled to 1 mm isotropic resolution using tri-cubic interpolation. The CSF and GM segmentation maps were rigidly registered to the corresponding FOD map and resampled to 1×1×1 mm^3^. The subject-to-template linear and nonlinear transformations from the group-wise registration were saved and used to warp and average all T1w and T2w scans to obtain the unbiased T1w and T2w brain templates. While the FOD template has a 1mm isotropic spatial resolution, the T1w and T2w brain templates were made available in a resolution of 0.5×0.5×0.5 mm^3^ for the whole brain and 0.3×0.3×0.3 mm^3^ for the central region that contains the subcortical structures, similar to the MNI-PD25 atlas.

With the resulting averaged FOD template, whole-brain tractography was performed using iFOD2 (Tournier et al., 2010) to produce an initial tractogram with 20 million streamlines, which was then filtered to 2 million streamlines with the SIFT (Smith et al., 2013) algorithm in MRtrix3 (Tournier, et al., 2019) to reduce false positives. To help with data visualization, a second, lighter version of the tractogram with 20K streamlines was also provide by randomly sampling streamlines from the 2 million-streamline tractogram.

### Subcortical segmentation and probabilistic tissue maps

The manual segmentations of 22 subcortical structures in the MNI-PD25 atlas (Xiao et al., 2019) were mapped to the resulting T1w population-averaged brain template by using T2w-to-T2w nonlinear registration (Xiao et al., 2019). The result was further revised manually by the author YX, who also performed the subcortical segmentation for the MNI-PD25 atlas. The segmented anatomical structures and their corresponding label numbers are listed in Table 1. By using the linear and nonlinear transformations from the group-wise registration in template making, the probabilistic tissue segmentation maps obtained from the FAST algorithm were warped and averaged in the T1w template space. The final population-averaged probabilistic tissue maps included white matter, cortical grey matter, subcortical grey matter, and CSF in the resolution of 0.5×0.5×0.5 mm^3^.

**Table 1:** Label numbers with the corresponding subcortical structures for subcortical segmentation of all atlases.

| Label number | Subcortical structure | Label number | Subcortical structure |
| --- | --- | --- | --- |
| 1 | Left red nucleus | 2 | Right red nucleus |
| 3 | Left substantia nigra | 4 | Right substantia nigra |
| 5 | Left subthalamic nucleus | 6 | Right subthalamic nucleus |
| 7 | Left caudate | 8 | Right caudate |
| 9 | Left putamen | 10 | Right putamen |
| 11 | Left globus pallidus externa | 12 | Right globus pallidus externa |
| 13 | Left globus pallidus interna | 14 | Right globus pallidus interna |
| 15 | Left thalamus | 16 | Right thalamus |
| 17 | Left hippocampus | 18 | Right hippocampus |
| 19 | Left nucleus accumbens | 20 | Right nucleus accumbens |
| 21 | Left amygdala | 22 | Right amygdala |

### Subthalamic nucleus parcellation

To facilitate the planning and post-operative analysis of deep brain stimulation surgery, the STN in the atlas was parcellated into sensorimotor, associative, and limbic sub-regions based on structural connectivity. With each subject’s FOD map, 10K streamlines were generated using the STN segmentation as the seeding mask for the sensorimotor, associative, and limbic regions defined from the Harvard-Oxford atlas (Desikan et al., 2006). Then, each functional sub-region was determined using a winner-takes-all approach with the normalized track density in each voxel within the STN segmentation at 1×1×1 mm^3^ resolution. Lastly, the individual STN parcellations were warped to the common template space at 0.3×0.3×0.3 mm^3^ resolution by using the previously obtained subject-to-atlas deformation fields, and the final population-averaged STN parcellation was generated based on a majority voting strategy based on all subjects. Note that the fiber tracking and functional region parcellation were performed for the left and right side separately.

## Ethics statements

All participants provided their written informed consent. The use of a publicly available, open dataset was approved by the University of Western Ontario Health Sciences Research Ethics Board (UWO HSREB) under protocol #108456 and by the Research Ethics Committee of Concordia University under protocol #30016659.

## CRediT author statement

*Please add a CRediT author statement for your data article here, using the <u>categories listed on this web</u>page*.

Yiming Xiao: Conceptualization, Methodology, Software, Data processing, Writing – Review & Editing

Jonathan C. Lau: Writing – Review & Editing

Terry Peters: Supervision, Writing – Review & Editing

Ali R Khan: Supervision, Data processing, Writing – Review & Editing

## Acknowledgments

The authors acknowledge the use of computer clusters of Digital Research Alliance of Canada for creating the brain templates.

